# Interresponse time and run length shaping by discrete-trial percentile schedule incorporating fixed consecutive number procedure

**DOI:** 10.1101/2024.05.07.592607

**Authors:** Tomotaka Orihara, Takayuki Tanno

## Abstract

The effect of incorporating a fixed consecutive number (FCN) procedure in interresponse time (IRT) shaping using a discrete-trial percentile schedule was examined. Eight pigeons were used. On each trial, pigeons were required to peck the IRT start key and then the IRT end key. Reinforcers were presented according to the percentile schedule based on the time interval (IRT) or the cumulative response count to the IRT start key during those two key pecks. Multiple responses to the IRT start key were allowed only in the condition incorporating the FCN procedure. Results showed that incorporating the FCN procedure improved the success of IRT shaping from 1/8 to 7/8 of the pigeons. The procedure developed in this study provides an ideal baseline for studying response shaping with IRT as the dependent variable.

## 1. Introduction

Response shaping is a method and phenomenon for acquiring new behavior (Pierce & Cheney, 2008). Differential reinforcement and successive approximation are the primary methods for the response shaping in non-human animals. For example, Ferguson and Rosales-Ruiz (2001) shaped horses to enter a trailer. They divided the shaping process into eight steps of successive approximation: differential reinforcement for approaching the trailer entrance, for placing parts of their body inside the trailer, for staying there for a certain duration, and so on. Since Skinner discovered response shaping in the 1940s (Peterson, 2004; Orihara & Tanno, 2022), it has been widely used in both experimental and applied areas.

Experimental analyses on the control variables of response shaping have been conducted using the percentile schedule (Platt, 1973). The percentile schedule is expressed by the following equation:

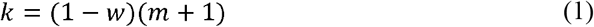

where *k* is the rank of the reinforcement criterion, *w* is the reinforcement probability, and *m* is the sample size. In this schedule, current response is reinforced or extinguished based on the *k*th response in the last *m* responses so that the reinforcement probability *w* is constant.

Dependent variable frequently used in studies on the percentile schedule was the interresponse time (IRT) in free-operant situation. Alleman and Platt (1973) manipulated the reinforcement probability, *w*, and showed that the lower *w*, the longer IRTs were shaped. Kuch and Platt (1976) also manipulated the reinforcement probability, *w*, by manipulating the duration of interreinforcement interval and showed that the longer the interval (the lower *w*), the longer IRTs were shaped.

Orihara & Tanno (2023) examined the IRT shaping in pigeon’s key pecking by percentile schedule in discrete-trial situation. At the beginning of each trial, either the left or the right key (IRT start key) turned on white. When the IRT start key was pecked, the key was turned off, and the other key (IRT end key) turned on white. When the IRT end key was pecked, the key was turned off, and the trial was finished. The IRT was defined as the time between the first and second responses. A food reinforcer was presented if the IRT meets the reinforcement criterion for the trial. Independent variables were the frequency of updates the reinforcement criterion and the presence or absence of reinforcement criterion retreat. In the former, they manipulated the update frequency of reinforcement criterion from pre response (it is a standard procedure of the percentile schedule) to per session. In the latter, while in the percentile schedule the IRT reinforcement criterion was shortened or lengthened based on the current performance (based on the last *m* IRTs), they prevented the shortening of the criterion. Their results showed that, while the latter independent variable strengthened the tendency of extinction, both of them seem to have no prominent effect on the accuracy and speed on IRT shaping.

Discrete-trial version of percentile schedule has two advantages over that of the free-operant version. First, it can distinguish changes in IRT length from changes in frequency of IRT occurrences. For example, suppose we want to shape a long IRT in a free-operant percentile schedule and we manipulate the reinforcement probability, *w*, as its independent variable. What is predicted is a decrease in the frequency of IRT occurrence under conditions where the reinforcement probability, *w*, is low. Then, it is not possible to distinguish whether the IRT became longer as a result of the long IRT shaping or whether the IRT became longer by decreasing the frequency of IRT occurrence due to low reinforcement probability. Conditions for shaping shorter IRTs must be considered to address this issue (Alleman & Platt, 1973). Discrete-trial version of percentile schedule allows to distinguish these two changes. The second advantage is the clarification of response unit for reinforcement. When a specific IRT is differentially reinforced in a free-operant situation, its effect may extend over a series of response sequences prior to reinforcement (Tanno & Silberberg, 2012; Tanno et al., 2015). In a discrete-trial procedure, it is possible to clearly define individual IRTs as response units.

One problem of Orihara and Tanno (2023) was that some pigeons failed to IRT shaping itself. Studies using percentile schedules in free-operant situations rarely find such results (Alleman and Platt, 1976; Kuch & Platt, 1976). Experimental analysis on controlling variables of response shaping needs stable baseline performance. Future studies using the discrete-trial version of percentile schedule should improve its procedure to allow for obtaining stable baseline performance of IRT shaping.

Orihara and Tanno (2023) proposed the fixed consecutive number (FCN) procedure (Mechner, 1958) as a candidate for improvement. Assume there are two types of responses, A and B. The FCN procedure is a procedure that presents a reinforcer for response B after *n* or more times of response A. The number of responses of response A before response B is called the run length. In a study by Machado and Rodrigues (2007) on pigeons, the run length was defined as the number of left key pecks before the right key pecks. They systematically manipulated the reinforcement criterion of run length from 4 to 32. As a result, the relative frequency distribution of run lengths showed a normal distribution centered around the reinforcement criterion. Galbicka et al. (2001) studied a schedule combining the FCN procedure with a percentile schedule in rat’s lever press. Here, the run length defined as the number of left lever presses before the right lever press. The target value of the run length was set at 16, resulting in an average of approximately 13 run length was shaped.

Why is it useful to incorporate FCN procedures into the Orihara and Tanno (2023) procedure? The shaping of long IRTs was attempted in this study. However, since reinforcers cannot be obtained without responding, short IRTs were sometimes shown by some pigeons, in the form of responding to the IRT start key at the start of the trial and pecking at the IRT end key shortly thereafter. The increase in such short IRTs, by the nature of the percentile schedule, precludes updating the reinforcement criteria in the direction of reinforcing longer IRTs. Such “impulsive” short IRTs have also been frequently observed under differential reinforcement of low rate schedules that reinforce longer IRTs (Kramer & Rilling, 1970). The FCN procedure would allow subjects to continue responding to the same key after the initial response to the IRT start key, thereby suppressing the occurrence of “impulsive” short IRTs.

Present study examined the incorporation of the FCN procedure into the Orihara and Tanno’s (2023) experiment to enhance the success of shaping long IRTs in a discrete-trial version of percentile schedule. Two experiments were conducted. In Experiment 1, we replicated Orihara and Tanno’s procedure to check the reproducibility of their results, and then examined whether the incorporation of FCN procedure would improve the results. The reinforcement criterion here was, accord with the FCN procedure, the cumulative number of responses to the IRT start key before responding to the IRT end key. In Experiment 2, the reinforcement criterion was changed to the time interval between the first response to the IRT start key and the response to the IRT end key. In this case, the number of responses to the IRT start key before responding to the IRT end key had no effect on presenting the reinforcer. In both experiments, both the run length and the IRT duration were analyzed as dependent variables.

We predicted the following results. In Experiment 1, the incorporation of the FCN procedure to Orihara & Tanno’s procedure should result in shaping long run lengths in the IRT start key. In this case, if we defined IRT as the time interval between the first response to the IRT start key and the first response to the IRT end key, then this should also be longer as well as the run length. In Experiment 2, while run length is irrelevant to the reinforcement criterion here, the response to the IRT start key continues to be made more than once, which lengthens the IRT. If these results are obtained, it reveals that the modification of the Orihara and Tanno (2023) procedure to keep the key light of the IRT start key on until the end of the trial after a single response to it is effective in shaping long IRTs.

## 2. Method

### 2.1 Subjects

Each of the four pigeons (*Columba livia*) was used for Experiments 1 and 2, respectively. Pigeons for Experiment 1 (MP1801, MP1803, MP1806, and MP1807) had an experimental history similar to the present experiment. Pigeons for Experiment 2 (MP1810, MP2106, MP2107, and MP2108) had an experimental history of stimulus equivalence of colors and shapes. Pigeons’ weights were maintained at approximately 80% of their free-feeding body weight. In their home cages, pigeons were free to access a water-tube, and were received mixed grains following each experiment sessions, if necessary. The pigeon’s room operated on a 12:12 h light/dark cycle, with lights turning on at 7:30 AM. Pigeon maintenance and experimental procedures were conducted in accordance with the Animal Experiment Committee of Meisei University.

### 2.2 Apparatus

For Experiment 1, four identical operant chambers with internal dimensions of 32.5 cm (length) × 32.5 cm (width) × 31.5 cm (height) were used. Except for the Plexiglas door and metal grid floor, all other surfaces were made of aluminum. The front panel contained three response keys (MED Associates, Inc., ENV-123AM) with a diameter of 2.5 cm, positioned horizontally 22.0 cm above the floor and 8.0 cm away from the center, and a feeder opening, which measured 6.2 cm in width and 5.4 cm in height, centered horizontally 5.7 cm above the floor. Only the left and the right keys were used, and those keys were transilluminated white, red, or green.

For Experiment 2, two identical operant chambers (MED Associates, Inc., ENV-007) with internal dimensions of 29.0 cm (length) × 24.0 cm (width) × 29.0 cm (height) were used. Except for the front and rear panels of the aluminum, and metal grid floor (MED Associates, Inc., ENV-005P), all other surfaces were made of Plexiglas. The front panel contained three response keys (MED Associates, Inc., ENV-126AM) with a diameter of 2.5 cm, positioned horizontally 22.0 cm above the floor and 9.0 cm away from the center, and the feeder opening, which measured 6.5 cm in width and 5.0 cm in height, centered horizontally 3.0 cm above the floor. Only the left and the right keys were used, and those keys were transilluminated white, red, or green.

A 28 V light bulb was used as the house light. Reinforcement consisted of access to hemp seeds for 3-s, delivered by a feeder (MED Associates, Inc., ENV-205M). The feeder light was illuminated during the reinforcer presentation. White noise masked extraneous sounds. A personal computer (Windows 7 Professional, Intel® Core™ i5-4690, 8 GB) with MED-PC IV software and a PC-compatible interface (Med Associates, Inc., DIG-716B) was used to control all experimental events and data recording.

### 2.3 Procedure

Present study consisted of two experiments that included common experimental conditions. We first describe those experimental conditions and then describe how they were combined in Experiments 1 and 2.

#### 2.3.1 General procedure and experimental conditions

##### General procedure

In each trial, pigeons were required to peck first at the left key (IRT start key) and peck last at the right key (IRT end key). The trial lasted for 50-s, with the house light turned on, followed by 10-s of inter-trial intervals (ITIs), with all lights turned off. If pigeons did not peck the IRT start key within 20-s from the start of the trial or did not peck the IRT end key within 20-s from the first peck to the IRT start key, key lights were turned off if they were illuminated. Each session consisted of 60 trials, conducted once daily, six days a week.

##### IRT reinforcement condition

The purpose of this condition was to replicate the difficulty of shaping long IRT lengths when using Orihara & Tanno’s (2023) procedure. In each trial, pigeons were required to peck once to the IRT start key and then once to the IRT end key. At the start of each trial, a key light on the IRT start key was turned on white. When the IRT start key was pecked, the key light was turned off, and a key light on the IRT end key was turned on white. Then the IRT end key was pecked, the key light was turned off. The IRT length in each trial was defined as the time between these two pecks. A reinforcer was presented if the IRT met the reinforcement criteria. Pecks to the not-illuminated key had no consequence.

##### Run reinforcement condition

At the start of each trial, a key light on the IRT start key was turned on red and a key light on the IRT end key was turned on green. Lights on both keys were turned off when one or more pecks to the IRT start key followed by a single peck to the IRT end key. Each peck followed by 0.1-s blinking the pecked key’s light. The run length was defined as the number of pecks to the IRT start key before a peck to the IRT end key, and the reinforcer was presented when this run length met the reinforcement criteria.

##### Modified IRT reinforcement condition

The procedure was the same as the run reinforcement condition except for the reinforcement criteria. In this condition, the IRT length was defined as the time between the first peck to the IRT start key and the subsequent peck to the IRT end key, and the reinforcer was presented when this IRT length met the reinforcement criteria.

#### 2.3.2 Experiment 1

##### Preliminary training

Because pigeons served for the Experiment 1 had experimental history of operant conditioning of key pecks, no shaping training was required. Before the run reinforcement condition, those pigeons were exposed to a mixed FCN fixed-ratio (FR) 1 schedule (Mechner, 1958). This schedule consisted of FR trials with proportion p and FCN trials with proportion 1-p. In FR trials, the reinforcer was presented for a single peck to the IRT start key (FR 1). In FCN trials, the reinforcer was presented for one or more pecks to the IRT start key followed by a single peck to the IRT end key. The p value was changed in the order of 0.50, 0.25, and 0.00; values were changed when a switching response from the IRT start key to the IRT end key was observed with a probability of more than 0.8 on the FCN schedule trial in the recent session. After the p value reached 0.00, this session was repeated three times.

##### Training

Pigeons were exposed first to the IRT reinforcement condition and then to the run reinforcement condition. A percentile schedule described as Eq. (1) was applied during the IRT reinforcement condition. Parameter values of the percentile schedule were equal to those of Galbicka et al. (1993), that is, *m* = 24, *w* = 0.33, and *k* = (24 + 1) (1 - 0.33) = 16.75. Consequently, IRT lengths from the last 24 trials were ordered from shortest to longest on an ordinal scale, and the IRT length ranked 17th on the ordinal scale was used as the reinforcement criterion for the next trial (the reinforcer was presented if the IRT length for the next trial was longer than that criterion). At the start of the second and subsequent sessions, the IRT of the last 24 trials of the previous session was carried over. Reinforcement criteria in the FCN reinforcement condition were determined in the same manner, except that run length, instead of IRT length, was used for reinforcement criterion. The IRT reinforcement condition was conducted for 20 sessions (but 10 sessions for MP1801) and the FCN reinforcement condition was conducted for 30 sessions.

A pre-condition phase preceded each condition. In this phase, pigeons were exposed to a schedule in which the reinforcer was presented for any switching response from the IRT start key to the IRT end key. This schedule continued for five sessions, and the phase was completed if the performance in its final session met the following criteria: For the IRT reinforcement condition, (1) IRTs were recorded in at least 48 out of 60 (80%) trials, (2) mean IRT length for those trials was less than 2-s, and (3) standard deviation of IRT lengths for those trials was less than 2; For the run reinforcement condition, (1) runs were recorded in at least 48 out of 60 (80%) trials, (2) mean run length for those trials was less than 3 responses, and (3) standard deviation of run lengths for those trials was less than 2. Pigeons that did not meet these criteria were then exposed, until they met them, to the following schedule:

For the IRT reinforcement condition, they were exposed to a schedule in which reinforcers were presented when the recorded IRT was less than 2-s; For the run reinforcement condition, they were exposed to a schedule in which reinforcers were presented when the recorded run was less than 3 responses.

#### 2.3.2 Experiment 2

##### Preliminary training

Before the IRT reinforcement condition, pigeons served for Experiment 2 were exposed to a preliminary training consisting of the following three steps. In the first step, to habituate the operant chamber, pigeons were left in the chamber with free access to food from the feeder. This training continued for three sessions, each lasted for at least 30 min. In the second step, pigeons were exposed to an autoshaping procedure (Brown & Jenkins, 1968). One of the three key lights was randomly turned on white for 10-s on variable time (VT) 30-s, and a reinforcer was presented immediately after the key light was turned off. The 12 intervals comprising each of the VT schedule were derived using the Fleshler and Hoffman (1962) progression and were randomly sampled without replacement. If pigeons pecked the illuminated key, the key light was turned off, and a reinforcer was presented immediately after that. Each session lasted for 60 reinforcer presentations. This autoshaping training continued until the reinforcer presentation was contingent on the pigeon’s key pecking for 48 out of 60 reinforcer presentations. In the third step, pigeons were exposed to a variable interval (VI) schedule in which a reinforcer was presented for the first response after the completion of an inter-reinforcement interval (IRI) that varies from one reinforcer to the next. A schedule was randomly assigned one of three keys during each IRI while the key light was turned on white. The VI value was increased from 10-s to 20-s to 30-s, each of which conducted for two, three, and four sessions, respectively. The 12 intervals comprising each of the VI schedule were derived using the Fleshler and Hoffman (1962) progression and were randomly sampled without replacement. Each session lasted for 60 reinforcer presentation or 60 min.

Before the modified IRT reinforcement condition, pigeons were exposed to a mixed FCN FR 1 schedule (Mechner, 1958). The procedure was the same as Experiment 1.

##### Training

Pigeons were exposed first to the IRT reinforcement condition and then to the modified IRT reinforcement condition. A percentile schedule described as Eq. (1) was applied in both conditions. Parameter values of the percentile schedule were *m* = 23, *w* = 0.33, and *k* = (23 + 1) (1 - 0.33) = 16. Consequently, IRT lengths from the last 23 trials were ordered from shortest to longest on an ordinal scale, and the IRT length that ranked 16th on the ordinal scale was used as the reinforcement criterion for the next trial (a reinforcer was presented if the IRT length for the next trial was longer than that criterion). At the start of the second and subsequent sessions, the IRT of the last 24 trials of the previous session was carried over. The IRT reinforcement condition was conducted for 20 sessions and the modified IRT reinforcement condition was conducted for 30 sessions. A pre-condition phase, which was the same as Experiment 1, was preceded for each condition. In other regards, the procedure was the same as Experiment 1.

## 3. Results and Discussion

We present a summary result of Experiments 1 and 2. Figure 1 shows the mean IRTs and mean run length for the last session of the pre-phase and each session of experimental conditions. The black circles in the first and third rows of Figure 1 shows the results of the IRT reinforcement condition, which conducted as a replication of Orihara & Tanno (2023). Their study had shown a failure of IRT shaping in some pigeons, and this result was reproduced here: The IRT was not lengthened in seven of the eight pigeons. Sessions for MP1803 was stopped after 10 sessions because this pigeon showed the mean IRT exceeded 10-s in the initial session and maintained it for 3 sessions.

**Figure 1.**
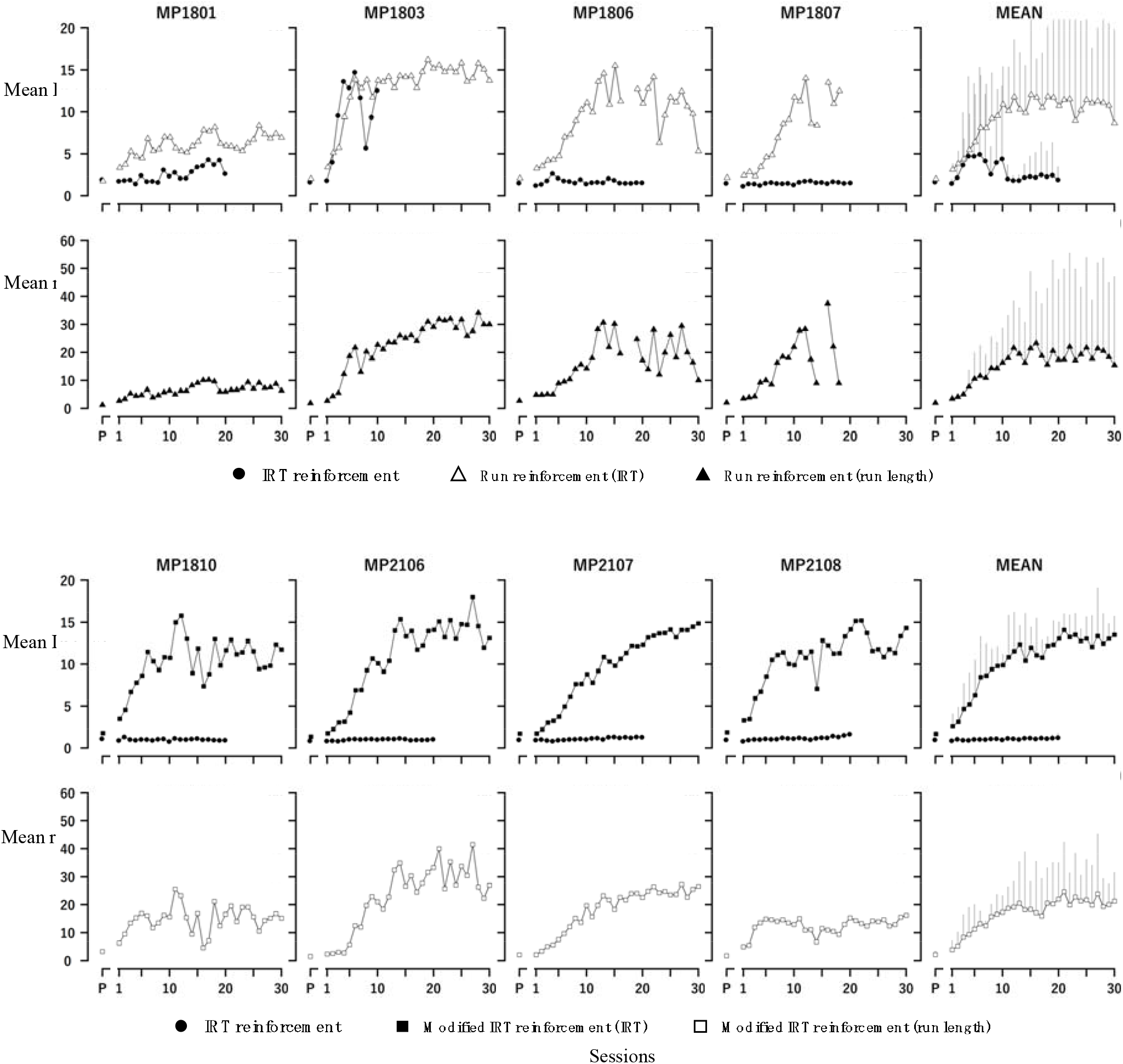
Mean IRTs and mean run length as a function of number of sessions for each experimental condition and subject in Experiments 1 and 2. *Note*. The error bars in the MEAN panel indicate 95 percent confidence intervals. On the X-axis, “P” represents the last sessions of the pre-condition phase, and the other numbers represent number of sessions in the condition phase. The upper limit of Y-axis for mean IRT is 20-s due to the experimental procedure.

Did the run reinforcement condition and the modified IRT reinforcement condition lengthened the IRT and the run length? The black triangles in the second row of Figure 1 shows mean run length in the run reinforcement condition in Experiment 1. Sessions for MP1807 was stopped after 22 sessions because this pigeon did not show a run (at least two pecks for the IRT start key) for consecutive three sessions. The mean run length was lengthened in three of the four pigeons. The white triangle in the first row of Figure 1 shows corresponding IRTs (converted from mean run length to the time interval between the first response to the IRT start key and the first response to the IRT end key). As the run length increased, so did these IRTs. The black squares in the third row of Figure 1 shows the mean IRTs in the modified IRT reinforcement condition in Experiment 2. IRTs were lengthened in all pigeons. The white squares in the fourth row shows the number of responses to the IRT start key (i.e., run length) between the first response to the IRT start key and the response to the IRT end key. As the IRTs lengthen, so did these run lengths. These results indicate that incorporating the FCN procedure into the Orihara & Tanno (2023) procedure is effective for lengthening IRTs.

Regression analysis between the IRT and the run length revealed close relationship between these two dependent variables. Upper and lower panels of Figure 2 show the relationship between the IRT and the run length for the run reinforcement condition of Experiment 1 and the modified IRT reinforcement condition of Experiment 2, respectively.

**Figure 2.**
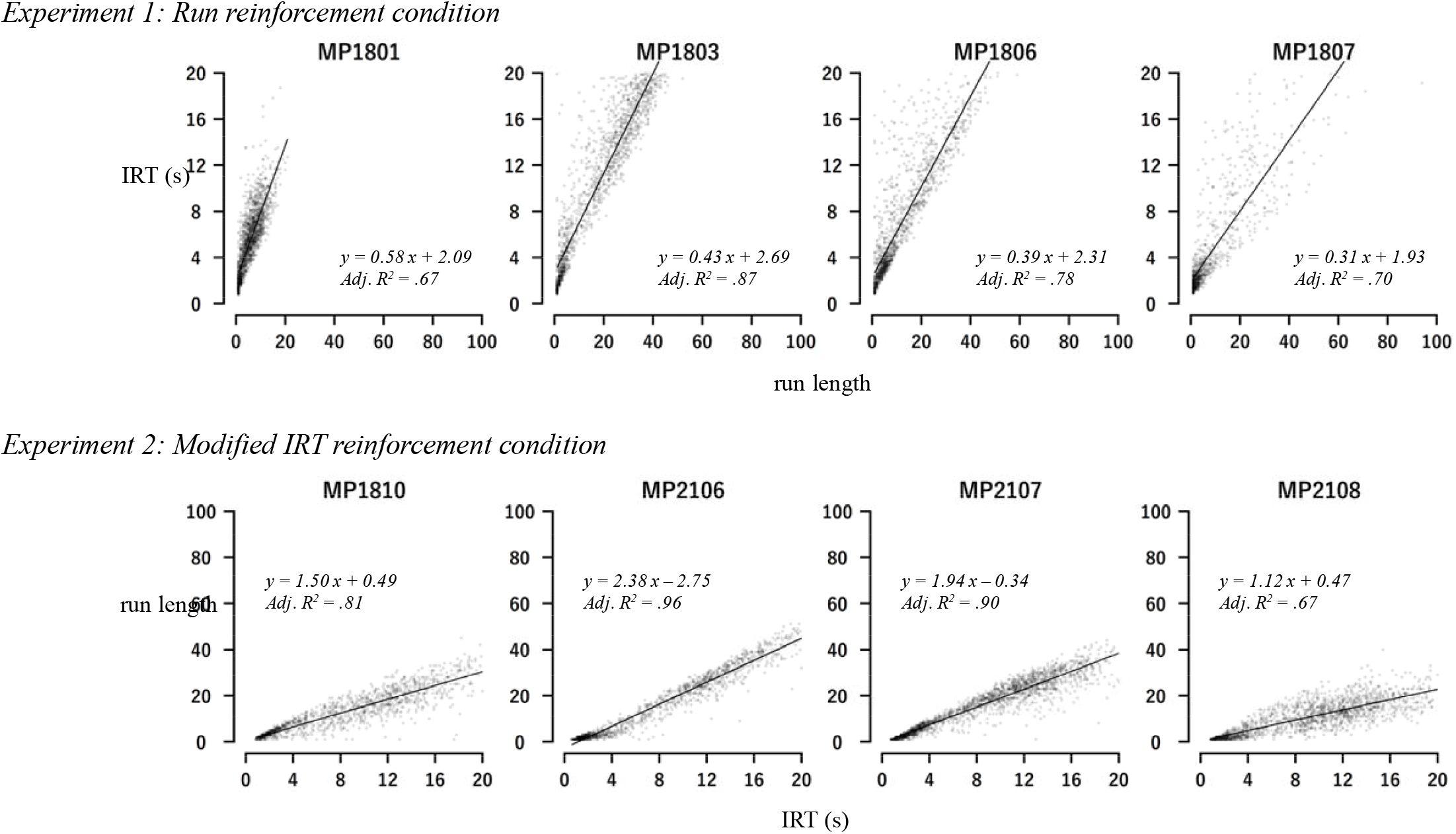
Top: Regression analysis for IRTs as a function of the run lengths in run reinforcement condition of Experiment1. Bottom: Regression analysis for run lengths as a function of the IRTs in modified IRT reinforcement condition of Experiment 2.

Since the reinforcement criterion in the upper panel was run length, the run length is plotted on the x-axis and IRT on y-axis. Similarly, the reinforcement criterion in the lower panel was IRT, so the x- and y-axes in the upper panel are reversed. Linear relationships between IRT and run length were observed in all pigeons. Regression analysis showed a statistically significant slope at p < .001 for all individuals, although the regression coefficient values varied. The coefficients of determination ranged from 0.67 to 0.96, demonstrating high explanatory power.

In the introduction, we argued that the failure of IRT shaping in Orihara & Tanno (2023) procedure was due to the impulsive IRTs. Figure 3 shows the relative frequency distribution of IRTs in Experiment 1. Row and column correspond to subject and five-session block, respectively. The results for the IRT reinforcement condition were as follows. For MP1806 and MP1807, the IRT distribution was narrow with a peak at 2-s, regardless of the session block. For MP1801, the IRT distribution had a peak at 2-s, but its range increased as the session progressed. For MP1803, the IRT distribution had no peak and was distributed from 2 to 20-s. In the run reinforcement condition, no pigeon showed the IRT distribution peaked at 2-s even in the first block. The IRT distributions shifted towards longer with session progression. In the sixth block, MP1801 and MP1806 showed a peak around 8 to 12-s, and MP1803 showed a peak at 16-s or longer. While sessions for MP1807 was stopped after 22 sessions, this pigeon showed a peak around 12-s at that point. The same analysis was conducted for the data of Experiment 2, and Figure 4 shows the results. The tendency observed in Figure 3 was reproduced in this Figure 4: the IRT distribution had a peak at 2-s in IRT reinforcement condition, but it disappeared in modified IRT reinforcement condition. These results indicate that “impulsive” short IRTs occurred in the IRT reinforcement condition, as was predicted, and incorporating the FCN procedures is effective to prevent such IRT occurrences.

**Figure 3.**
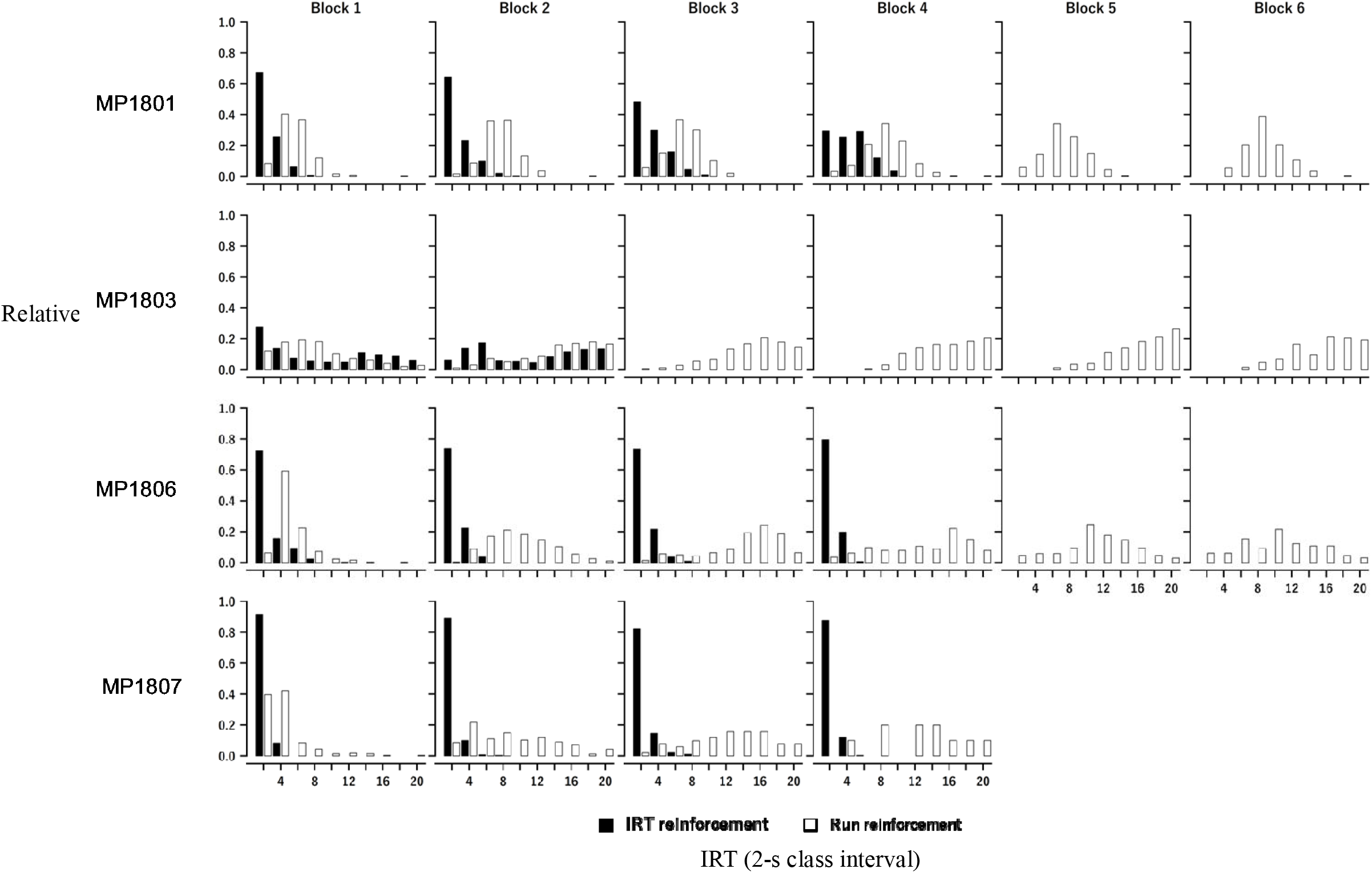
Relative frequency distribution of IRTs in IRT reinforcement condition and run reinforcement condition of Experiment 1.

**Figure 4.**
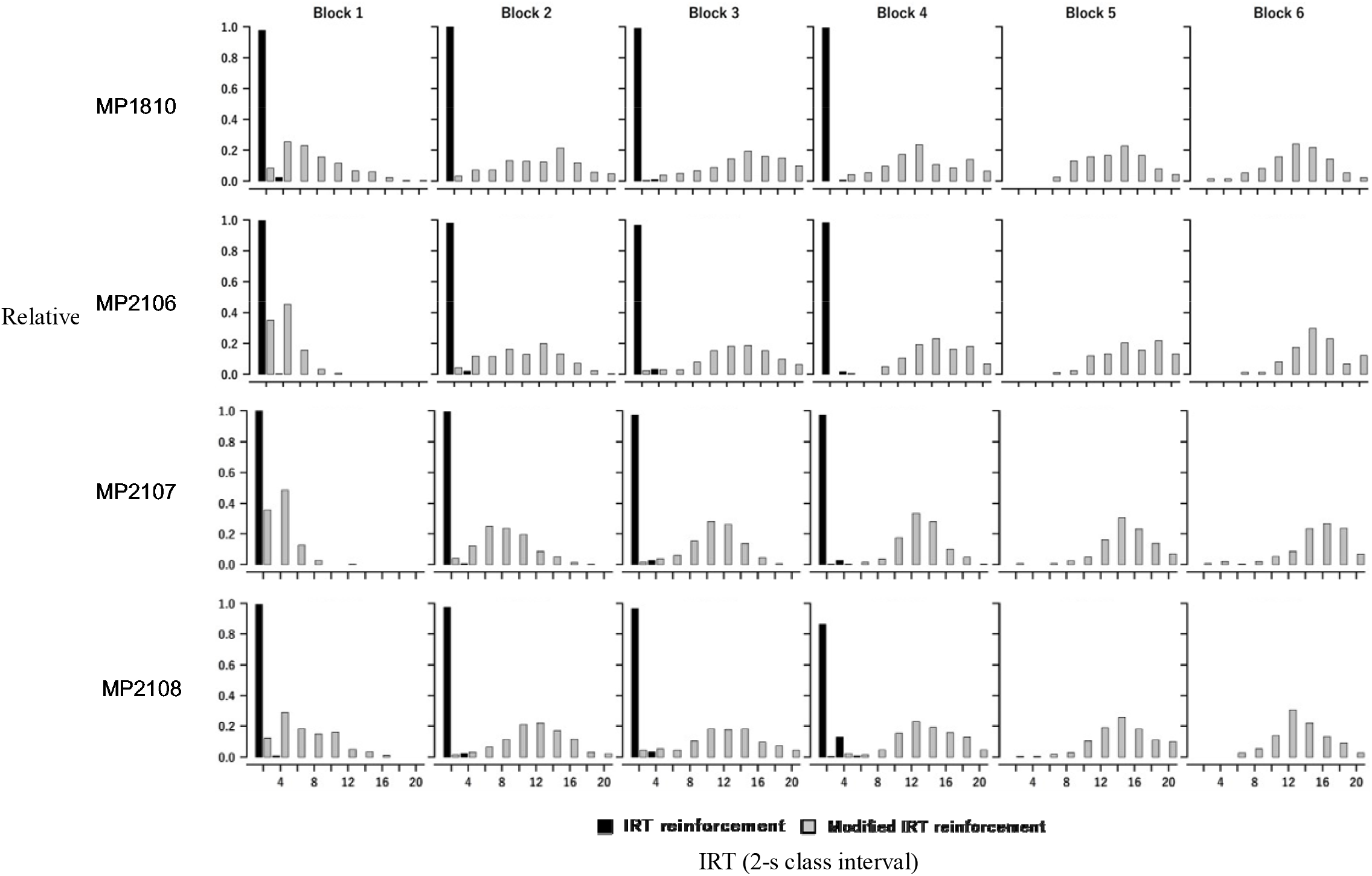
Relative frequency distribution of IRTs in IRT reinforcement condition and modified IRT reinforcement condition of Experiment 2.

Table 1 shows probability of reinforcement and probability of IRT occurrence for each experimental condition of Experiments 1 and 2. The probability of reinforcement is calculated, for each session, as dividing the number of reinforcer presentations by the number of trials with IRT occurrence. Values in Table 1 are average across all sessions for each experimental condition for each subject. The values were approximately 0.33 in all condition and subject, which equal to the expected value of reinforcement probability when w = .33 in Eq. (1). The IRT occurrence is calculated, for each session, as dividing the number of trials with IRT occurrence by the total number of trials. Values in Table 1 are average across all sessions for each experimental condition for each subject. In Experiment 1, three pigeons (MP1803, MP1806, MP1807) showed a large difference in probability of IRT occurrence between conditions, and on average, the value was .32 lower (a decrease of approximately 19 times of IRT occurrences for 60 trials) in run reinforcement condition than in IRT reinforcement condition. In Experiment 2, three pigeons (MP1810, MP2106, MP2108) showed a large difference in probability of IRT occurrence between conditions, and on average, the value was .28 lower (a decrease of approximately 17 times of IRT occurrences for 60 trials) in modified IRT reinforcement condition than in IRT reinforcement condition. One possible reason for the decrease in the probability of IRT occurrence is the time limit on response opportunities. In this experiment, there was a 20-second time limit between the first response to the IRT start key and the response to the IRT end key. In conditions where the IRT or run length was successfully lengthened, the trial ended before the response to the IRT end key so the response opportunity was lost, which may have reduced the IRT occurrence probability.

**Table 1.** Probability of reinforcement and probability of IRT occurrence for each experimental condition of Experiments 1 and 2.

| Measure | Condition | Experiment 1 |  |  |  |  | Experiment 2 |  |  |  |  |
| --- | --- | --- | --- | --- | --- | --- | --- | --- | --- | --- | --- |
|  |  | MP1801 | MP1803 | MP1806 | MP1807 | MEAN | MP1810 | MP2106 | MP2107 | MP2108 | MEAN |
| **Probability of reinforcement** | IRT reinforcement | 0.32 | 0.39 | 0.32 | 0.33 | 0.34 | 0.35 | 0.35 | 0.37 | 0.37 | 0.36 |
|  | Run reinforcement | 0.28 | 0.32 | 0.28 | 0.32 | 0.30 |  |  |  |  |  |
|  | Modified IRT reinforcement |  |  |  |  |  | 0.29 | 0.32 | 0.37 | 0.36 | 0.34 |
| **Probability of IRT occurrence** | IRT reinforcement | 1.00 | 0.90 | 1.00 | 1.00 | 0.98 | 0.99 | 1.00 | 1.00 | 0.99 | 1.00 |
|  | Run reinforcement | 0.86 | 0.65 | 0.59 | 0.54 | 0.66 |  |  |  |  |  |
|  | Modified IRT reinforcement |  |  |  |  |  | 0.56 | 0.62 | 0.93 | 0.77 | 0.72 |

The results are summarized as follows. To improve the shaping of longer IRTs in discrete-trial version of the percentile schedules, we tested the incorporation of the FCN procedure into the Orihara and Tanno’s (2023) procedure. The results were positive. While the original Orihara and Tanno’s procedure (IRT reinforcement condition) showed successful IRT shaping only in 1/8 pigeons, this was improved to 7/8 by incorporating the FCN procedure (Figures 1). This improvement can be attributed to the fact that the incorporation of the FCN procedures prevented “impulsive” short IRTs (Figures 3 & 4).

Another feature of the improvement is the strong correspondence between IRT and run length (Figure 2). Is the success of IRT shaping simply a byproduct of shaping the run length? This question requires a conceptual discussion of IRT. The IRT is defined as a response property that can be changed by reinforcement contingencies (Morse, 1966).

Pigeons engage in other behaviors (e.g., grooming) during the time interval measured as IRT (Iversen, 1991; Palya, 1992). Differential reinforcement of IRTs can be considered to be the differential reinforcement of such behavior patterns as a whole (Tanno & Silberberg, 2012; Tanno et al., 2015). In other words, IRT is a measure of response patterns that include other behaviors, and in this study, run length is another measure of such other behaviors. If so, the correspondence between run length and IRT (Figure 2) is a necessary consequence.

Experimental analyses on control variables for response shaping have been conducted with IRT as a dependent variable while manipulating percentile schedule variables as independent variables (Alleman & Platt, 1973; Galbicka & Platt, 1986; Kuch & Platt, 1976). A baseline of IRT shaping is needed if the accuracy and speed of the IRT shaping are to be compared in quantitative manner. The changes in IRT in Figure 1 demonstrated a typical learning curve. We believe that the procedures reported by this study provide an ideal baseline for experimental investigations of reaction formation using percentile schedules.

